# Application of mechanistic multiparameter optimization and large scale in vitro to *in vivo* pharmacokinetics correlations to small molecule therapeutic projects

**DOI:** 10.1101/2024.03.06.583780

**Authors:** Fabio Broccatelli, Vijayabhaskar Veeravalli, Daniel Cashion, Javier L. Baylon, Franco Lombardo, Lei Jia

## Abstract

Computational chemistry and machine learning are used in drug discovery to predict target-specific and pharmacokinetic properties of molecules. Multiparameter optimization (MPO) functions are used to summarize multiple properties into a single score, aiding compound prioritization. However, over-reliance on subjective MPO functions risks reinforcing human bias. Mechanistic modeling approaches based on physiological relevance can be adapted to meet different potential key objectives of the project (e.g. minimizing dose, maximizing safety margins and/or minimized drug-drug interaction risk) while retaining the same underlying model structure. The current work incorporates recent approaches to predict *in vivo* PK properties and validates *in vitro* to *in vivo* correlation analysis to support mechanistic PK MPO. Examples of use and impact in small molecule drug discovery projects are provided. Overall, the mechanistic MPO identifies 83% of the compounds considered as short-listed for clinical experiments in the top 2^nd^ percentile, and 100% in the top 10^th^ percentile, resulting in an area under the receiver operating characteristic curve (AUCROC) > 0.95. In addition, the MPO score successfully recapitulates the chronological progression of the optimization process across different scaffolds. Finally, the MPO scores for compounds characterized in pharmacokinetics experiments are markedly higher compared to the rest of the compounds synthesized, highlighting the potential of this tool to reduce the reliance on *in vivo* testing for compound screening.

## INTRODUCTION

Predictive modeling-driven small molecule drug discovery relies on computational chemistry and machine learning (ML) approaches to predict target-specific and pharmacokinetic (PK) properties of new molecular entities (NME).^1^ In addition, physiologically based PK (PBPK) approaches are utilized to mechanistically predict absorption, distribution, metabolism, elimination (ADME), therapeutic dose, and drug-drug interaction (DDI) potential of NMEs.^2^

Multiparameter optimization (MPO) functions are used in compound progression and design to summarize multiple *in vitro* and *in silico* properties (e.g. potency, clearance, permeability, solubility) into a single score.^3^ Without computer-aided analysis, it is practically impossible to readily and accurately balance more than four variables.^4^ While the benefits of reducing complexity to inform decision making with MPO functions are evident, over-reliance on subjective MPO functions may ultimately reinforce human bias.^5^

Accurate predictions of *in vivo* PK and pharmacodynamic (PD) properties are required to inform clinical candidate selection. Although the approaches to predict *in vivo* PK/PD properties of drug candidates may vary from user to user, there is a consensus in the industry around the use of mechanistic modeling approaches.^6^ Unlike empirical MPO functions, the mechanistic approaches are based on physiological relevance, can be tested, and translated across species, and can support estimation of safety margins and DDI risk. In addition, the interpretability of mechanistic models is a further key differentiation from “black box” methods; drug designers can leverage the model structure and the observed *in vitro* to *in vivo* correlations to identify opportunities to design better molecules. In this work, we demonstrate the value of using mechanistic PK models as an MPO scoring function for compound testing and progression ^7-9^. We incorporate and extensively validate recent approaches to the prediction of *in vivo* PK properties, provide adequate examples of *in vitro* to *in vivo* correlation (IVIVc) analysis in support of MPO, and illustrate some examples of use and impact in small molecule drug discovery projects.

## METHODS

### PKPD Assumptions

The choice of MPO method depends on the known or assumed relationship between PK and PD (PK/PD). Quantitative PK/PD models are valuable during clinical candidate selection for dose projection and DDI assessment, especially for targets and modalities (such as covalent drugs or targeted protein degraders) resulting in delayed responses (indirect response models).^10^ In early phases, PK/PD relationship can be simplified at the expense of accuracy to fit into three response categories: C_max_, C_min_, and C_avg_ driven efficacy. When PK/PD relationships are simplified as C_max_ driven, the objective of dose optimization is to achieve a transient exposure higher than the minimum efficacious concentration (MEC). In this case, high potency, high bioavailability (F), fast absorption, moderate clearance (CL), and short half-life (T_1/2_) are desirable. However, C_max_ is less commonly associated with on-target activities and often associated with off-target and safety-related concerns. When PK/PD relationships are simplified as C_min_ driven efficacy, the efficacious dose is determined based on the terminal concentration at steady state, matching the MEC. In this case high potency, high F, low CL, and prolonged oral T_1/2_ are desirable, which is a reasonable default assumption for competitive inhibitors. Lastly, with PK/PD relationships simplified as C_avg_ driven efficacy, the efficacious dose is determined based on the average concentration at steady state (that is, exposure over-dosing interval) matching the MEC. For C_avg_ driven efficacy, high potency, high F, and low CL are desirable, while T_1/2_ is believed to be less important. Notably, a C_avg_ that is at least 5-fold higher than the total EC_50_ predicts efficacious dose for over 90% of the 164 drugs studied by Jansson-Löfmark et. al.^11^ This assumption can be used as a reference for early projects for which information about the PKPD driver is not available and subsequently updated when more quantitative information about the PKPD drivers becomes available.

### Prediction of PK

All the non-proprietary datasets used for model validation and calibration in this work are available as supporting information. This study used physiologically based methods to predict PK parameters. The hepatic well-stirred model predicts total metabolic CL from *in vitro* measured or *in silico* predicted CLint,app.^12^ CL is assumed to be predominantly mediated by hepatic metabolism. The free drug hypothesis is leveraged to derive CL_int_ from CL_int,app_ and fraction unbound in microsomes (equation 1):

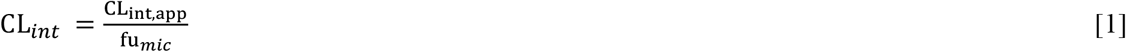

Blood to plasma partitioning is assumed to be unity for all compounds in this study, and hepatic blood clearance is derived using equation 2, where fu_p_ is the fraction unbound in plasma and Q is the physiological liver blood flow (20.7 mL/min/Kg in human)^12^:

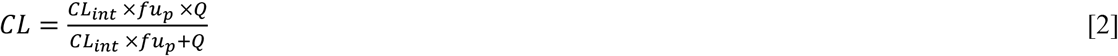

Volume of distribution (VD_ss_) at steady state is predicted using the Korzekwa-Nagar model (equation 3), which characterizes distribution in three compartments: plasma, tissue water, and tissue lipids.^13, 14^ R1 is the concentration ratio of plasma proteins in tissue water-to-plasma, estimated to be between 0.052 and 0.116. An average value of 0.084 is adopted in this study. V_p_ and V_t_ are the volume of plasma (0.043 L/Kg in human) and tissue (0.557 L/Kg in human), respectively. LK_L_ (product of lipid concentration and association constant for drug binding to lipids) can be predicted based on the fraction unbound in microsome (fu_mic_) at 1 mg/mL microsomal protein concentration (equation 4)

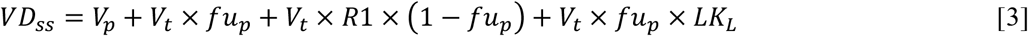

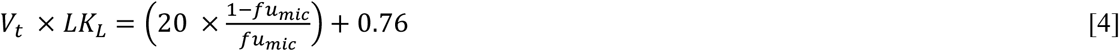

Equation 3 can be re-arranged to estimate fu_p_ from *in vivo* VD_ss_ and measured fu_mic_ as follows:

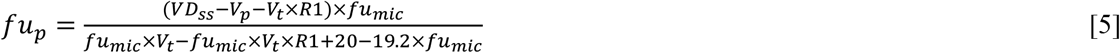

The use of this back-calculated fu_p_ in IVIVc is disclosed for the first time in this work.

MRT (mean residence time of a molecule in the body following an IV dose) and its inverse, k_e_ (rate of elimination), can be estimated from CL and VD_ss_ using equations 6 and 7:

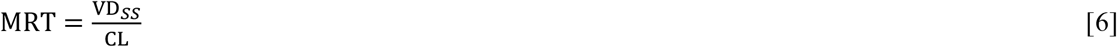

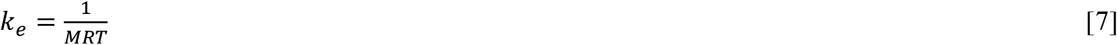

CL predicted leveraging the well-stirred model is also used to estimate the fraction surviving first pass elimination in the liver (F_h_) following oral absorption as a fraction of the liver blood flow using the following equation:

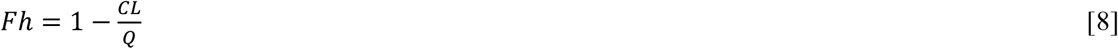

This assumption is valid for compounds that are primarily eliminated via hepatic metabolism. Fraction surviving first pass elimination in the gut following oral absorption (F_g_) is assumed to be unity due to the lack of predictive *in vitro* and *in silico* tools to predict this parameter.

The fraction absorbed in the intestine (F_a_) is calculated using the model presented by Yu, which estimates the absorbable dose (D_abs_) based on the physiological parameters such as intestinal transit time (Ti, 3.3 hr adopted for human) and intestinal surface area (Si, 800cm^2^ adopted for human), as well as the compound-specific parameters, including solubility and effective jejunum permeability (P_eff_) as shown in equation 9.^15, 16^ F_a_ can be calculated using equation 10, which is a ratio of D_abs_-to-administered dose. The Fa value should be capped to a maximum of 1:

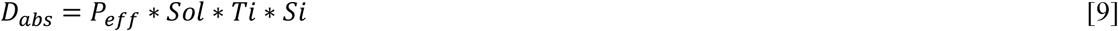

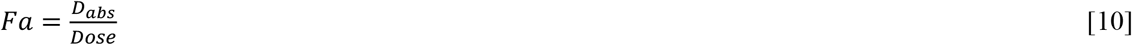

The use of intestinal solubility for the estimation of D_abs_ is a reasonable assumption for neutral, basic, zwitterionic and anionic compounds, for which the stomach solubility do not significantly exceed the intestinal solubility. Weakly basic compounds are typically well solubilized in the stomach due to the low gastric pH and can exhibit a markedly lower solubility in the intestine due to the prevalence of their neutral form. When the rate of precipitation from supersaturated solution is slow for a weakly basic compound, this has the potential to remain in solution during the intestinal transit time. Rate of precipitation is not a simple property to model, so an empirical approach is adopted to model and simplify this phenomenon. A D_abs_ value based on solubility in the stomach is calculated, and the final adopted D_abs_ value is based on intestinal solubility when this is higher than stomach solubility. The average of D_abs_ based on intestinal and stomach solubilities is adopted in all the other cases to simulate an intermediate level of compound precipitation. A human dose of 210 mg is used in all simulations (3 mg/kg).

The human P_eff_ value is estimated from MDCK apparent permeability (MDCK P_app_) data based on a linear fitting of the data published by Varma et al (equation 11), resulting in a R^2^ of 0.65 based on Log permeability of 17 marketed drugs (Figure 5). ^17^

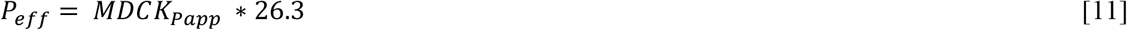

The literature on predicting rate constant for absorption (k_a_) with simple approaches is limited. In this work we leverage the Sinko model^18^:

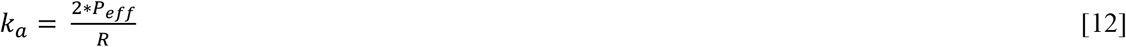

In this equation, the R stands for radius of human intestine, which is 1.75 cm. ^19^ In order to avoid unrealistic k_a_ values, this is capped at a lower boundary of 0.2 hr and a higher boundary of 1 hr. This model provides an optimistic estimate of k_a_ given the lack of consideration for dissolution; however, the model could be further evolved by leveraging mechanistic PBPK approaches. ^20^ Finally, oral bioavailability (F) can be derived from the following equation:

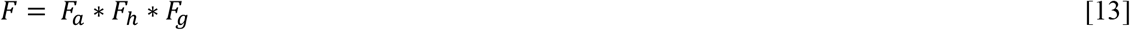

Oral PK curves are generated using a one-compartment distribution model, chosen for its simplicity.^7^ The average, maximum, and minimum concentration of the PK curve at steady state are defined by equations 14, 15, and 16, respectively:

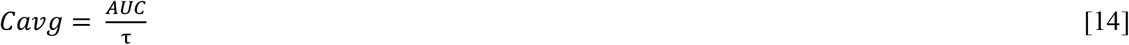

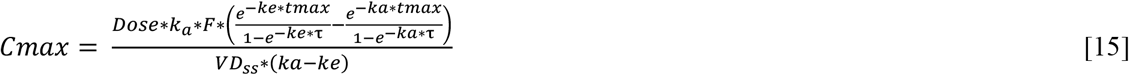

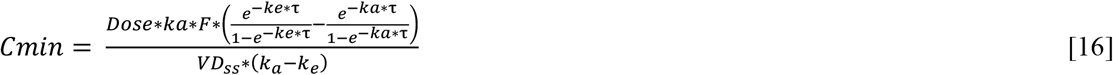

Where *τ* is the dose interval, t is time, t_max_ is the time corresponding to C_max_, calculated as:

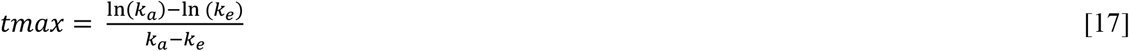

AUC is the area under the simulated PK curve and can be calculated by the following equation:

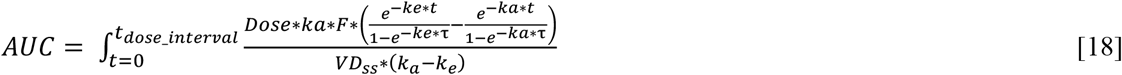

The coverage scores for average, maximum, and minimum concentrations (C_avg_, C_max_, C_min_), can be derived using the following equations:

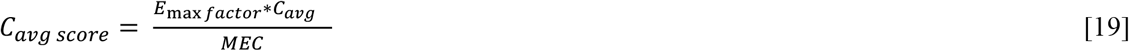

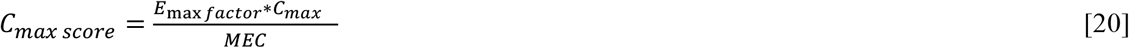

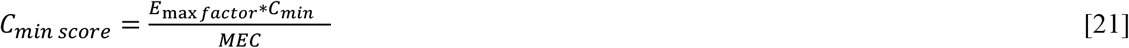

Here, minimum efficacious concentration (MEC) represents the total *in vivo* potency, which is estimated from apparent *in vitro* intrinsic potency (EC_50,app_) and binding estimates in potency media (fu_media_), as well as fu_p_. This is calculated using equation 22 assuming same free concentration in plasma and at the site of action.

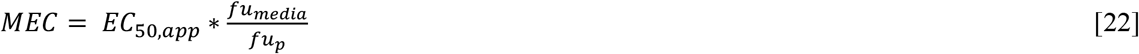

When measurements in the fu_media_ are not available, this can be estimated based on the Fu_p_ in full plasma and by leveraging the dilution formula noted below, where D is the dilution factor for protein content in media-to-plasma (e.g. D=10 for a 10-fold dilution) ^21^:

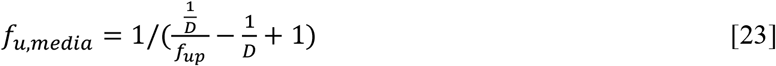

The application of this equation to the estimation of f_u,media_ makes the reasonable assumption of similarity between the behavior of compounds toward human (or animal) plasma proteins and FBS (fetal bovine serum) typically used in cellular screens.

The E_max factor_ is an adjustment factor to consider the depth of *in vitro* bioactivity. This is defined as:

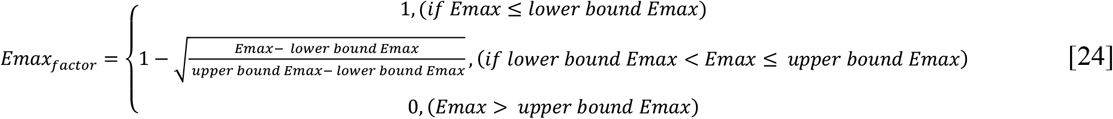

The lower bound E_max_ is a value below which we believe the depth of degradation is thorough. The upper bound E_max_ is a value above which we believe the depth of degradation does not bear degradation activity for effective coverage score calculation.

The method may also include an off-target coverage score. For example, a hERG score can be defined as the following:

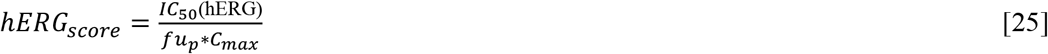

Where IC_50_ hERG is the hERG inhibition potency in a dose response assay.

The off-target score can be combined with the coverage score of choice to provide a surrogate therapeutic index score (TI score) which is defined as the following, assuming the off target in this case is hERG inhibition:

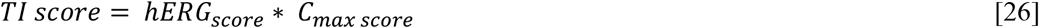

Input values from six experimental assays (fu_mic_, fu_p_, CL_int,app_ in microsomes, kinetic solubility at pH 7.4, potency, and MDCK P_app_) were derived from experimental assays where possible. If these values were not available, proprietary machine learning models were used instead^3^. Additionally, predicted pK_a_ values were utilized to estimate solubility at pH 2 (stomach pH) and pH 7.4 (intestinal pH). Solubility at different pH is calculated leveraging Henderson and Hasselbach equation as:

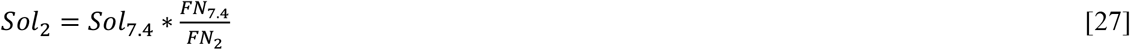

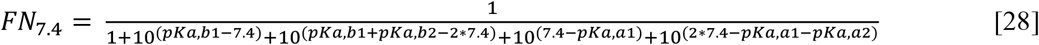

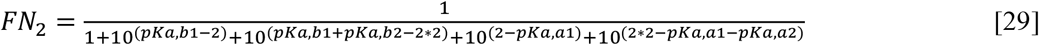

Where FN is fraction neutral. pK_a,a_ and pK_a,b_ are the most acidic and basic pK_a_, respectively; pK_a,a2_ and pK_a,b2_ are the second most acidic and basic pK_a_, respectively.^22^ When no acidic or basic pK_a_ are available, use 14 for acidic and 0 for basic.

A prediction of LK_L_ based on physicochemical properties is developed using the same dataset and following the same method as described by Berellini et al ^23^:

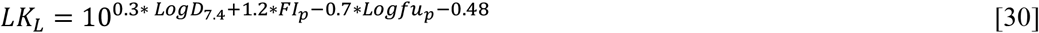

Where L is the amount of lipid in the tissue, and K_L_ is the association constant for drug binding to the lipid. FI_p_ is the fraction positively ionized, a value that can be derived from equation 28 by setting the pK_a,a1_ and pK_a,a2_ values to 14.

Equation 30 and equation 4 can additionally be combined to predict fu_mic_ from physico-chemical properties (equation 31).

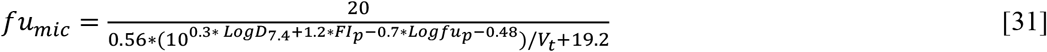

The accuracy in predicting CL, fu_mic_ and VD_ss_ is quantified as the absolute average fold error:

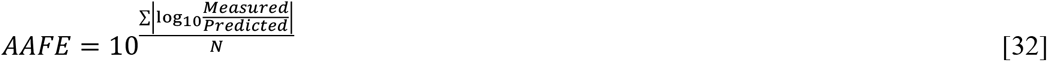

The area under curve of the receiver operating characteristic (AUCROC) curve is used to measure accuracy of ranking and is calculated using the pROC package.^24^

## RESULTS AND DISCUSSION

### In vitro to in vivo correlations of CL

The pre-requisite for the effective use of a mechanistic MPO model is the existence of solid IVIVc within the scaffold of interest; IVIVc disconnects for PK properties may point at additional elimination/distribution mechanisms that in turn rely on ancillary *in vitro* and *in silico* models to adequately describe these mechanisms. As a default assumption, discovery projects typically assume phase 1 Cytochrome P450 mediated metabolism as the primary mechanism of elimination, which is experimentally tested with NADPH-supplemented microsomal stability assays. When scaffold IVIVc disconnects are observed between microsomal stability and *in vivo* pre-clinical data, this assumption can be revised leveraging follow-up studies using hepatocytes, liver S9 fractions, bile duct canulated PK, PK in the presence of chemical inhibitors or metabolite identification.

In addition, accuracy of fu_p_ measurements should also be evaluated. Plasma protein binding assays are prone to provide inaccurate estimates for compounds that have chemical or plasma stability issues, for compounds that are highly non-specifically bound to the assay apparatus, and for compounds that, due to the high extent of binding, fail to reach equilibrium under standard assay conditions.^25^ For advanced compounds, customized measurements (e.g. extended incubation time, diluted plasma, bi-directional dialysis) are typically performed to solidify the fu_p_ estimates. One parsimonious approach to understanding the cause of CL IVIVc disconnects is to leverage the prediction of VD_ss_ from fu_mic_ to back-calculate an *in vivo* fu_p_ (equation 5). Because VD_ss_ and CL are independent PK parameters, the back-calculated *in vivo* fu_p_ value can subsequently be used to estimate *in vivo* CL. In project 1 (Figure 1, top graph), characterized by highly bound compounds with varying degrees of plasma instability, the initial method of choice for fu_p_ measurement was ultracentrifugation,^26^ which resulted in poor CL IVIVc correlation. However, when back-calculated *in vivo* fu_p_ was used instead, the correlation improved significantly (from negative correlation to R^2^ of 0.71 for LogCL).

**Figure 1.**
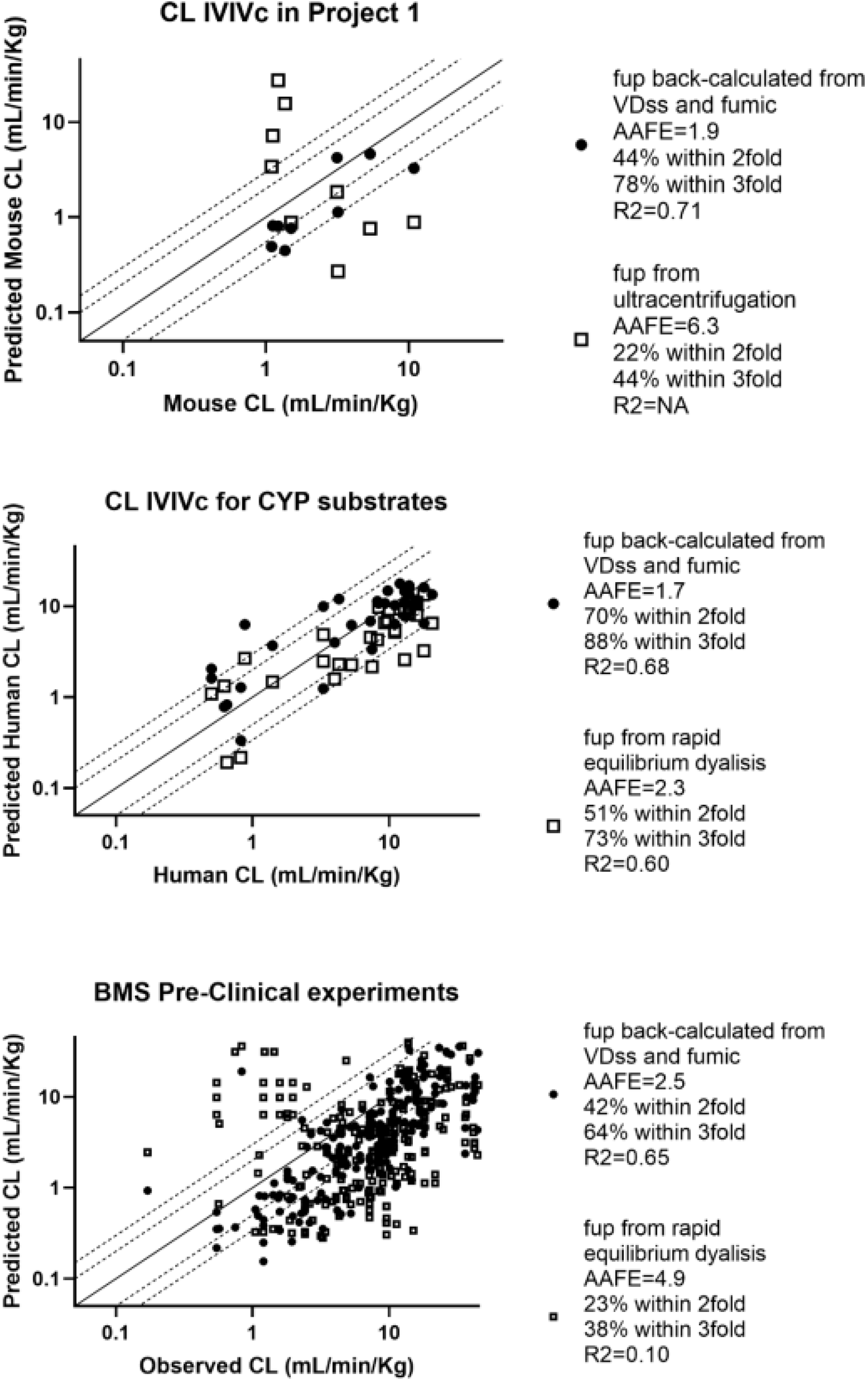
Clearance predictions across 3 different datasets. Empty squares represent predictions generated leveraging in vitro plasma protein binding data. Full circles represent predictions generated leveraging plasma protein binding data back-calculated from *in vivo* volume of distribution at steady state leveraging the Korzekwa-Nagar method. Accuracy for the different approaches is quantified as absolute average fold error (AAFE), R^2^, as well as percentage of predictions within two and three-fold.

These observations are not unique to Project 1. Jones et al. recently presented a dataset of *in vitro* and *in vivo* data to assess CL predictions based on human hepatocytes, in which a general underprediction of CL was observed.^27^ This was followed by a publication from the same group describing a new approach to measuring fu_p_, which significantly improved the CL prediction for a set of well-behaved Cytochrome P450 substrates for which CL IVIVc disconnects were previously observed.^28^ Figure 1 (middle graph) shows the extended original dataset presented by Jones et al., including 33 Cytochrome P450 substrates with measured CL_int_; all compounds included in this data exhibit CL_int_ values well within the dynamic range of the assay (>7 ml/min/kg). In this ideal dataset of well-behaved marketed drugs, the use of back-calculated *in vivo* fu_p_ measurements improved the absolute average fold error from 2.3 to 1.7, and the percent of predictions within 2-fold from 51% to 70%.

Figure 1 (bottom graph) additionally represents a similar analysis across all in-house compounds tested in mouse, rat, dog, or cynomolgus monkey. Since the mechanism of elimination for compounds in this dataset is unknown, property-based filters were adopted to focus on a chemical space in which the microsomal stability assay is likely to be representative of mechanisms of elimination, in agreement with previous reports about the chemical properties of P450 substrates (CACO2 P_app_ >5 × 10^−6^ cm/s, predicted LogD_7.4_>1, MW between 250 and 750, neutral or basic compounds).^17, 27, 29^ In this dataset, inclusive of numerous different scaffolds, a significant improvement in CL prediction accuracy is observed when the back-calculated *in vivo* fu_p_ was used: the absolute average fold deviation (AAFE) decreased from 4.9 to 2.5, R^2^ increased from 0.1 to 0.65, % within 2-fold increased from 23% to 42%. The accuracy in prediction can be expected to decrease when the dataset is expanded to include compounds for which the route of *in vivo* elimination is not well represented by the *in vitro* assay utilized or beyond the quantification limit of the assay. While the approach of using back-calculated *in vivo* fu_p_ appeared to improve accuracy in CL prediction, it is important to stress that this approach has value in retrospective analysis to understand IVIVc disconnects and can’t be used prospectively since CL and VD_ss_ are collected in the same experiment.

Based on these findings, it is recommended that fu_p_ measurements are investigated as a primary source of IVIVc CL disconnects by leveraging equation 5; when the re-analysis offers better correlation for *in vivo* and *in vitro* derived CL within the scaffold, the MPO functions presented in this work can be utilized to rank-order compounds in congeneric series. As previously discussed, fu_p_ has little influence on the T_1/2_ (that is, MRT) of neutral and basic compounds that are predominantly distributed in lipids; in addition, Liu et al discussed how the fu_p_ has comparable weightage on total CL and MEC, effectively canceling out in the C_avg_ based dose projection formula.^30^ A notable exception is represented by hydrophilic and negatively charged compounds, for which the primary distribution compartment is not in the lipids. For these molecules, fu_p_ values have a substantial influence on the predicted MRT, hence when C_max_ or C_min_ driven efficacy is assumed for these molecules, accurate estimates of fu_p_ might be required. Sensitivity analysis based on equations 2 and 3 can be utilized to assess the impact of fu_p_ on MRT in a specific congeneric series of molecules.

### *In vitro* to *in vivo* correlations of VD_ss_ and microsomal binding predictions

Lombardo et al. demonstrated that the mechanistic model described by Oie-Tozer is superior to allometric approaches in predicting human VD_ss_ across a large dataset of marketed drugs.^31^ The Oie-Tozer model leverages similar assumptions and structure as the recent model presented by Korzekwa and Nagar, with the former describing tissue affinity in terms of fraction unbound in tissues (fu_t_) and the latter using concentration and affinity to lipids (LK_L_).^14^ Berellini and Lombardo used a large set of marketed drugs with experimentally determined pKa and lipophilicity to derive a prediction of fu_t_ and VD_ss_ based on physico-chemical parameters.^23^ The accuracy of human VD_ss_ predictions based on physicochemical parameters is superior or comparable to the accuracy of predictions resulting from any individual pre-clinical species. While predictions based on LogD_7.4_ and pKa are of great value in the optic of model-driven drug discovery, these assays are not always available (especially pKa) during early phases for many compounds. Reliance on predicted values or surrogate chromatographic assays might produce significant deviation from the ideal values, as demonstrated by Chan et al. ^32^

In this work, the same dataset, method and physiological values presented by Berellini et al. is used to compare the accuracy of the Oie-Tozer (state of the art) and more recent Korzekwa-Nagar models.^13,14,23,31^ In the context of early dose projection, the value proposition for the Korzekwa-Nagar model is the ability to predict VD_ss_ from fu_mic_ values, a parameter that in lead optimization is frequently measured in parallel with microsomal CLint shortly after synthesis. Experimental VD_ss_ in preclinical species can additionally be used to estimate *in vivo* pre-clinical LK_L_, and subsequentially human VD_ss_.

Finally, a recent dataset with available experimental fu_mic_, predicted pK_a_, and chromatographic LogD_7.4_ is used in combination with the dataset presented by Lombardo et al as further validation (Set 2).^31, 33^ Results from this analysis highlighted similar accuracy between the two models; predictions based on *in vitro* data inputs emerge as a valid alternative to predictions based on early rodent PK screening for estimating human VD_ss_. Minor differences in the compound count reflects the exclusion from the Oie-Tozer predictions of compounds with aberrant fu_t_ as described by Lombardo et al.^31^ These results are summarized in Figure 2.

**Figure 2.**
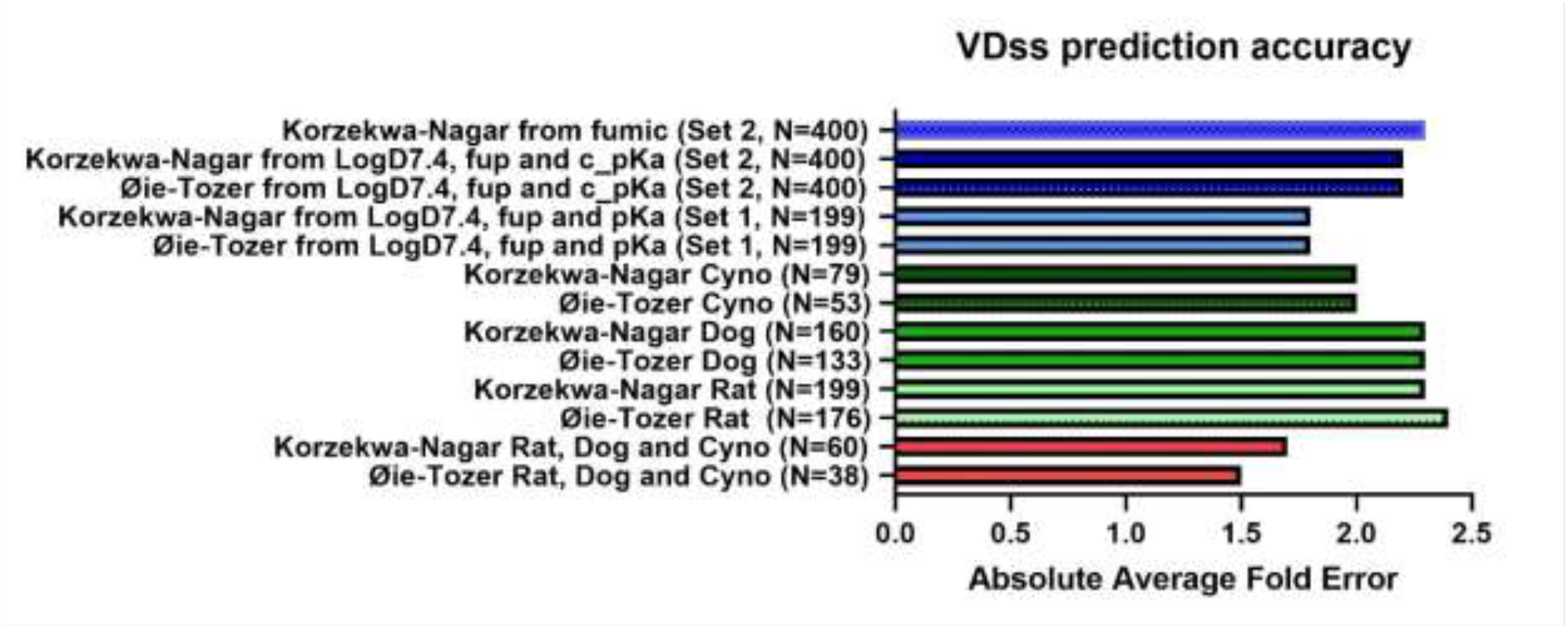
Predictions of Volume of distribution at steady state across different datasets and methods. Absolute average fold error (AAFE) values for the Oie-Tozer method are reported as presented in the reference publication by Lombardo et al.^31^

Winiwarter et al. reviewed the accuracy of different methods for predicting fu_mic_ from physicochemical properties and found that all the models reviewed demonstrated higher accuracy for compounds with cLogP<3, while predictions for more highly bound/lipophilic compounds were generally less accurate.^34^ In this work, the Turner model, which previously demonstrated state-of-the-art performance, is selected as a comparison for fu_mic_ predictions based on the combination of equations 30 and 4, leveraging the dataset presented by Tess et al.^33,34^ This is shown in Figure 3:

**Figure 3.**
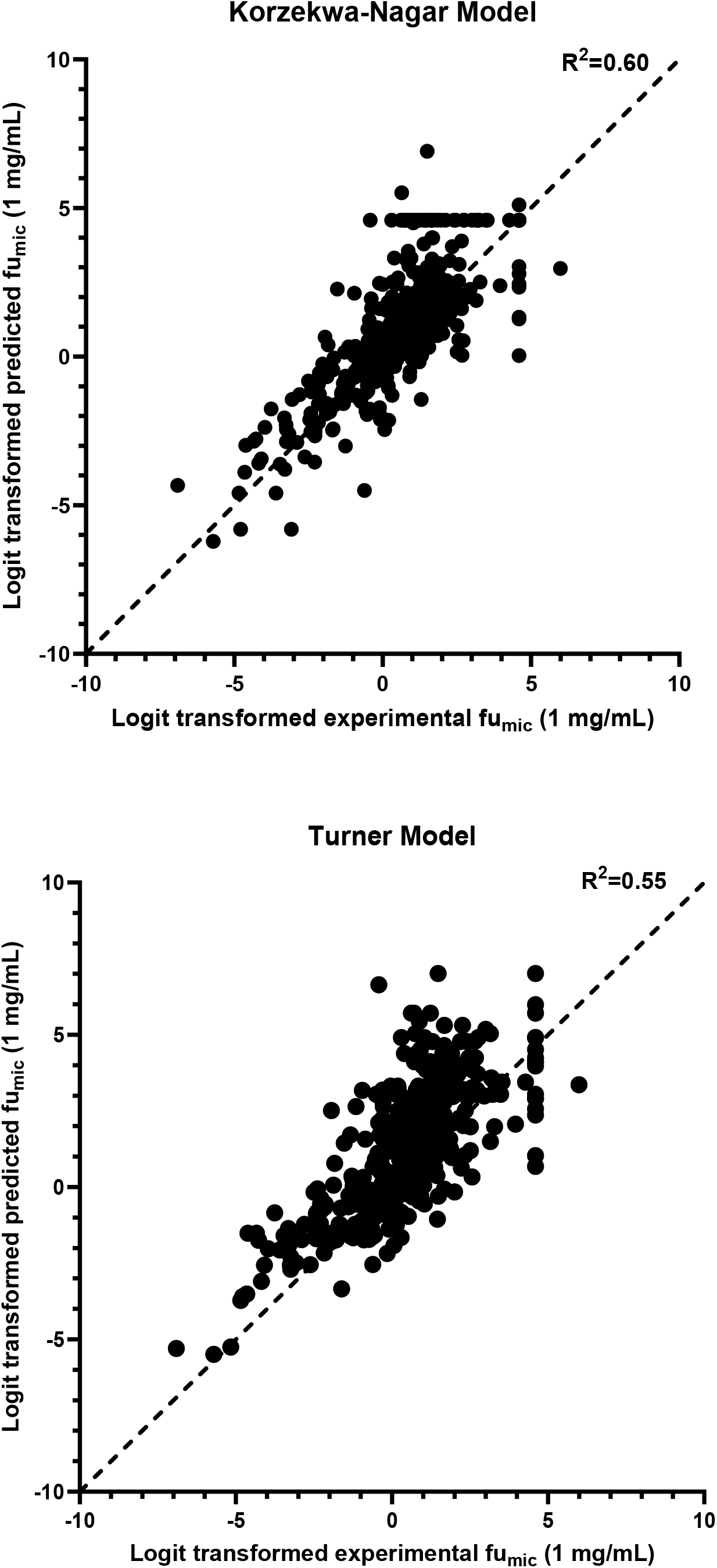
Predictions of microsomal binding based on physico-chemical properties. Experimental and predicted fu_mic_ were logit transformed logit(fu_mic_) = Ln(fu_mic_ /(1-fu_mic_)).

In the MPO application to BMS projects, machine learning machine learning models were used to estimate fu_mic_ due to their superior accuracy in the scaffold of interest.^34^ The predictions based on the equation 31 (presented in this work) is provided to allow the application of the MPO in chemical scaffolds and discovery projects for which accurate machine learning models are not readily available. Predictions of fu_mic_ based on equation 31 demonstrate superior performance compared with state-of-the-art models, while retaining self-consistence with VD_ss_ estimate and physico-chemical properties. In the internal BMS projects, fu_mic_ was always used as input for VDss predictions due to data availability (shake-flask LogD7.4 measurements were available for a limited number of compounds, while fu_mic_ measurements were regularly performed).

### *In vitro* to *in vivo* correlations of MRT

Cecere et al. previously characterized accuracy in MRT predictions from *in vitro* properties leveraging lipophilic metabolic efficiency (LipMetE).^35^ In this manuscript, “Set 2” is further refined to only include compounds that are primarily eliminated via metabolism (<20% excreted unchanged) and in a physicochemical space in which transporters are expected to have minimal influence on total CL (neutral or basic, P_app_ in MDCK > 5×10^−6^ cm/s and LogD_7.4_ > 1).^17, 27, 29^ For the prediction of VD_ss_, the Korzekwa-Nagar model based on physicochemical data is used when LogD_7.4_ < 3 (due to the high fu_mic_ values in this space making the estimate of LK_L_ from microsomes less quantitative); predictions based on the same approach as well as fu_mic_ were averaged for the remaining compounds. Figure 4 summarizes the comparative performance of the methods presented in this work and the method presented by Cecere et al. While quantitative predictions of MRT remain challenging, the methods presented in this work demonstrate good ability to differentiate shorter vs longer MRT compounds.

**Figure 4.**
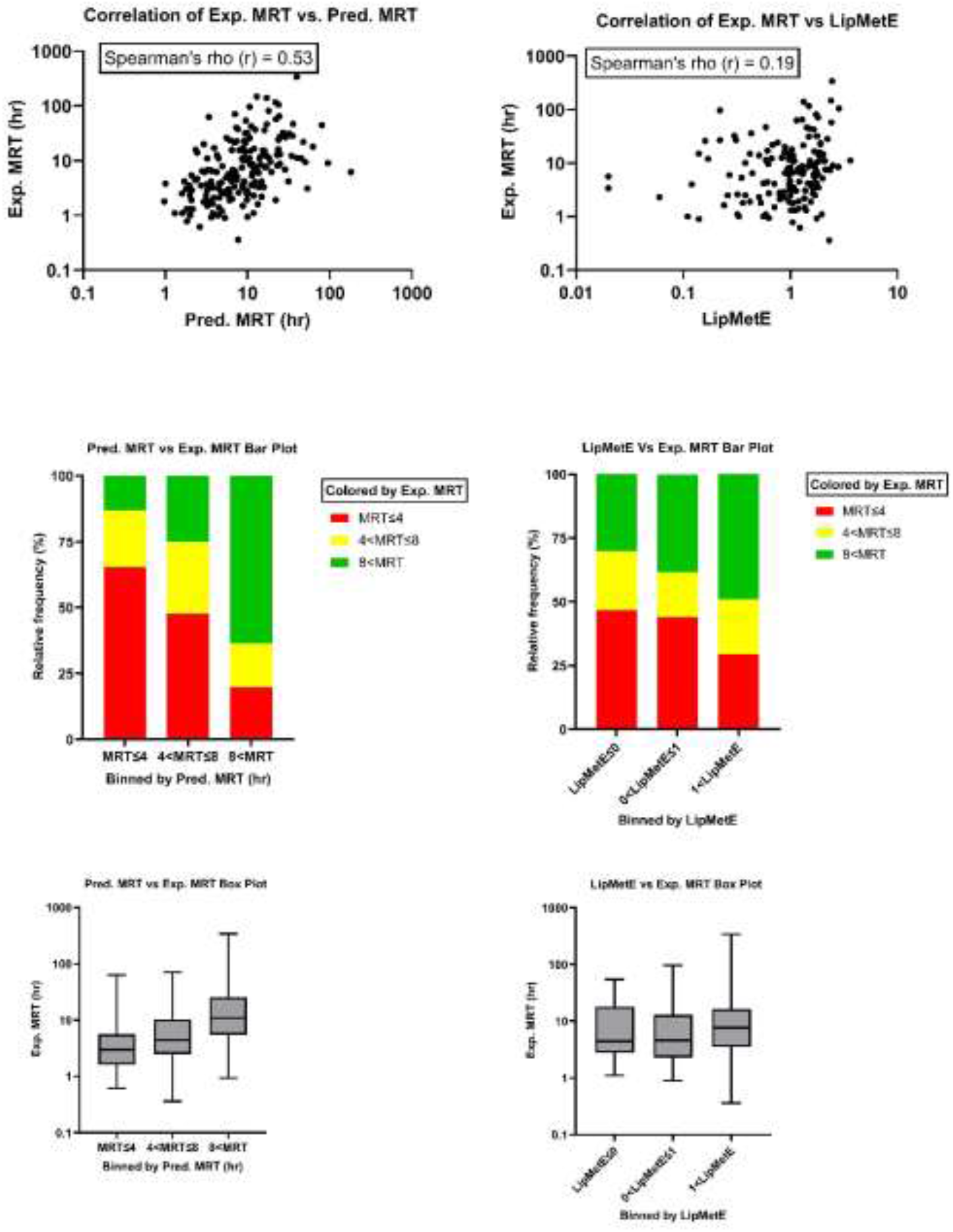
Comparison between intravenous mean residence time predictions in human from the combination of well-stirred hepatic model and Korzekwa-Nagar model, and lipophilic metabolic efficiency (LipMetE). Left side shows correlation between predicted MRT and experimentally measured MRT for assessing accuracy of MRT prediction. Right side shows correlation between calculated LipMetE and experimentally measured MRT for assessing the predictive power of LipMetE. Top pair of figures show scattered plot with Spearman’s rho ranking accuracy. Middle pair of figures illustrate a three-bin classification of predicted MRT and LipMetE, with experimental MRT and its relative frequency color-coded to each bin. Bottom pair of figures show box plot for three-bin classification of predicted MRT and LipMetE, with relative distribution of experimental MRT in each bin.

**Figure 5.**
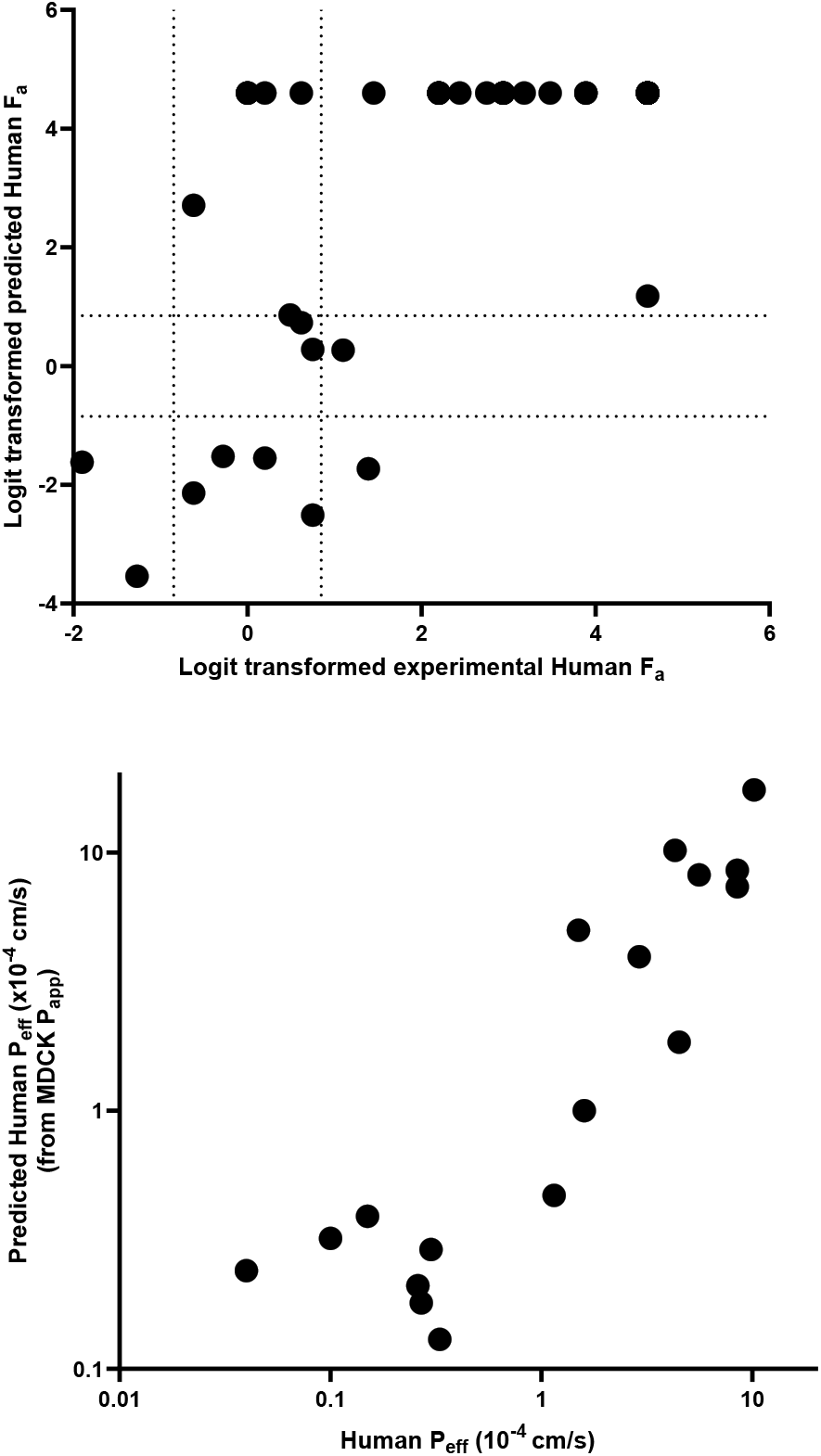
Prediction of fraction absorbed orally in human and human jejunum effective permeability from *in vitro* parameters. Experimental and predicted F_a_ were logit transformed logit(F_a_) = Ln(F_a_/(1-F_a_)), values equal to 1 were capped to 0.99 to avoid exclusion from the plot. Dashed lines correspond to F_a_ values of 0.3 and 0.7, defining low and high absorption categories.

The predicted MRT shows a reasonable rank correlation with experimental MRT (Spearman Rho = 0.53), while LipMetE exhibited a weaker correlation (Spearman Rho = 0.19). In addition, employing the LipMetE approach revealed approximately 30% of high MRT (MRT > 8 h) and low MRT (MRT < 4 h) compounds being misrepresented in the lower and higher bins for LipMetE, respectively (Figure 4). However, a significant improvement was observed when utilizing the predicted MRT from the Korzekwa-Nagar model, with approximately 13% and 20% of high MRT and low MRT compounds being incorrectly categorized into lower and higher bins for predicted MRT, respectively.

An additional benefit of using the Korzekwa-Nagar approach in conjunction with the well stirred model to predict MRT stems from the similarity in the input parameters for the two methods (fu_p_, fu_mic_). The error propagation emerging from experimental inaccuracies of fu_p_ and fu_mic_ measurements is reduced in the MRT estimate, compared to estimates of VD_ss_ and CL. For example, a compound with CL_int,app_ of 50 mL/min/kg and fu_mic_ equal to 0.9, variations in fu_p_ of up to 5-fold from the “true value” of 0.1, results in an approximately 5-fold variability in CL and/or VD_ss_, while such error imparts < 1.5-fold variability to the calculated MRT.

### *In vitro* to *in vivo* correlation of oral absorption

Oral absorption is largely a function of compound’s physico-chemical properties, passive permeability, and formulation; however, the majority of these parameters are unknown during early stages of research, which is a limitation for IVIVc based on pre-clinical data as well as for preliminary human dose projections; consequently, a qualitative assessment of F is a reasonable expectation before candidate nomination. Figure 5 presents the predictivity of the absorption models discussed in this study, using data published by Varma et al.^17^ The average error in predicting F_a_ for a set of 46 oral drugs is 0.15 (15% of % Fa). Notably, some compounds with low molecular weight, which have been previously associated with paracellular transport (absorption between tight junctions), are included in this set and incorrectly predicted as having low F_a_ (such as cimetidine, ranitidine, and methyldopa).^36^ Although this study does not explore thorough differentiation of absorption mechanisms, we attribute these discrepancies to the inability of the *in vitro* systems to account for paracellular transport accurately. Overall the correlation between the estimated and measured F_a_ was comparable to what reported by Varma et al (R^2^ or 0.43 vs 0.38).^17^

### Application of PK MPO function in small molecule drug discovery projects

This work provides examples of two small molecule projects, one of which assumes C_min_ driven efficacy (Project 1) and one assuming Cavg driven efficacy (Project 2). Compounds are divided in different cohorts of approximately same size (early stage, middle stage and late stage) based on registration dates. Because both project teams started from sub-optimal chemical matter and progressed to clinical candidates, the average MPO score is expected to improve over time to reflect the progress made during the lead optimization campaign.

Project 1 involved 571 compounds for which lack of selectivity did not limit progression; when different entries are associated with racemic mixture and enantiomeric pure compounds, only racemates are included in the analysis. 24 compounds were tested in rodent (4% of total) and 4 molecules in this project were short-listed as potential clinical candidates (0.7% of total). Figure 6 illustrates the MPO score for compounds in different cohorts. The project team focused on two scores: C_min_ score and a TI score (estimating hERG margins based on C_min_ score and C_max_ hERG score, Eq 26).

**Figure 6.**
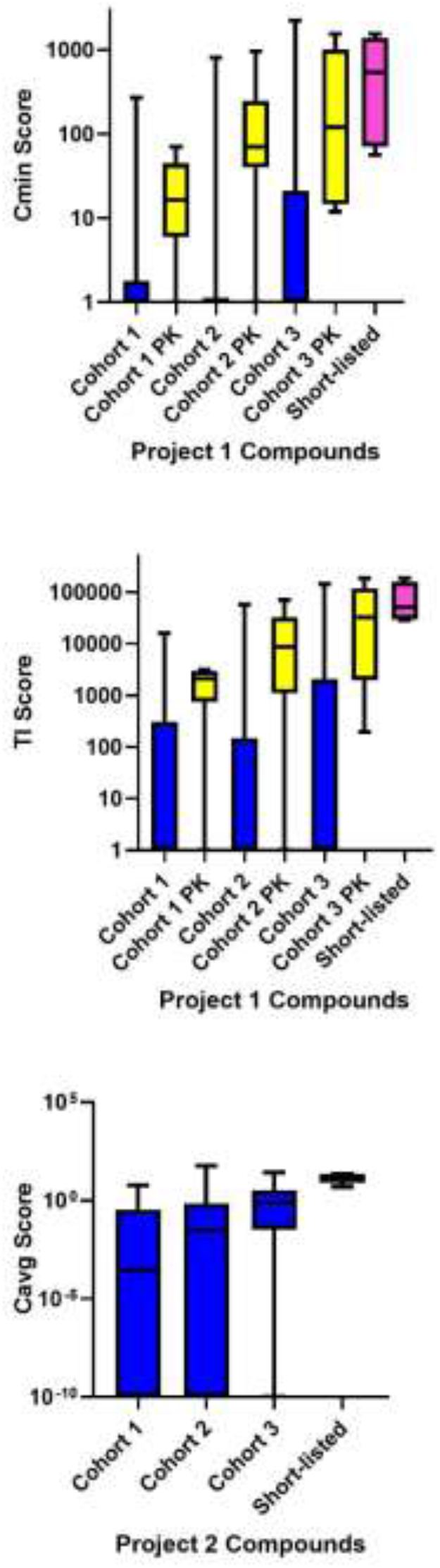
Application of the mechanistic pharmacokinetic multi-parameter optimization score in two internal projects. The different cohorts of compounds synthesized are based on the registration date; within the same cohort, the suffix PK indicates compounds tested *in vivo* in rodent, while the groups lacking the PK suffix are compounds that have not been tested *in vivo*. Short-listed compounds were considered as potential clinical candidates during the final stages of the lead optimization.

The PK MPO successfully parallels the chronological progression of the chemical series and the better quality of the compounds selected for *in vivo* studies, as well as those considered for clinical experiments. The group of 15 top ranked molecules based on the TI Score includes the four short-listed compounds, and 8 compounds that had been characterized in PK studies; overall the short-listed compounds were found in the top 2^nd^ percentile of the total distribution based on the TI score (AUCROC = 0.99). The prediction of compounds characterized *in vivo* based on the TI score results in an AUCROC = 0.86. The PK MPO score was developed in the context of project 1 and has been increasingly adopted starting from cohort 3 to identify compounds as candidates for *in vivo* PK and PK/PD experiments. The chemistry team became familiar with this method while approaching candidate selection. Although the method was not used to identify additional candidates this was leveraged during subsequent efforts to identify back-up compounds, leading to a significant reduction of *in vivo* testing.

Project 2 is a legacy campaign for which fewer details are available on PK and PK/PD experiments due to different informatics systems for compound experiment tracking. A total of 415 compounds are available in this set, of which 2 were short-listed as clinical candidates (0.4% of total). PK/PD experiments run at the early stages of the project indicated AUC driven efficacy, consequently, compounds were ranked based on their C_avg_ score. Similar to what was observed in Project 1, the score successfully recapitulates compound progression and identifies the short-listed lead compounds with a large margin compared to the distribution of the compounds synthesized in the campaign (Figure 6): one lead is in the 1^st^ percentile (5^th^ out of 415), while the other is in the 9^th^ percentile (39^th^ out of 415). Overall, the prediction of short-listed compounds based on the C_avg_ score results in an AUCROC = 0.95. The mechanistic PK MPO can be compared to more traditional linear scoring functions to gain insights on its differentiation and usefulness.^3^ In project 1, the linear MPO score used by the same team identifies the 4 short-listed compounds in top 30, but none of them ranks in the top 15. In project 2, both short-listed compounds are predicted between the 3^rd^ and the 8^th^ percentile. In short, the two methods offer orthogonal and complementary approaches to score compounds, the observed differences can inform additional strategies to compound optimization that may have not been initially envisioned by the team.

After the success observed in Projects 1 and 2, the mechanistic PK MPO method was applied to six additional ongoing internal projects, consistently identifying the top lead compounds within the top 10^th^ percentile of the scoring molecules. Currently, the PK MPO score is primarily used to rank-order synthesized compounds and inform *in vitro* and *in vivo* testing. Efforts to leverage the mechanistic PK MPO score during compound design are promising, but it is too early to draw meaningful conclusions.

## CONCLUSION

In this study, we introduce a methodology based on mechanistic PK science to rank compounds in small molecule therapeutic projects. The equations, methods, and datasets used in the mechanistic score are all available either in this manuscript or in the supporting information. The aim of this work is to produce high-throughput calculations to be incorporated in standard data mining software used by medicinal chemists for SAR; therefore, while we strived to leverage state of the art PBPK approaches for PK parameter prediction, no attempt was made to incorporate slower methods based on ordinary differential equations to describe the distribution of the test compounds across different organs.

Our retrospective IVIVc VD_ss_ analysis demonstrates that the Korzekwa-Nagar method exhibits accuracy comparable to the state of the art Oie-Tozer model.^31^ In addition, we further expand the scope of the Korzekwa-Nagar to derive *in vivo* distribution and microsome binding predictions from physio-chemical properties (pK_a_, fu_p_, and LogD_7.4_). Interestingly, predictions of microsome binding, indirectly obtained from the Korzekwa-Nagar method, perform similarly to previously published state-of-the-art methods for the same endpoint. Furthermore, we demonstrate how the *in vivo* fu_p_ back-calculated using the Korzekwa-Nagar method can be leveraged to understand IVIVc CL disconnects across multiple datasets. These findings highlight plasma protein binding assays as an underappreciated potential source of IVIVc disconnects, in agreement with previous observations from DMPK scientists in academia and industry.^25-28, 33, 37-39^ The Korzekwa-Nagar model combined with the well-stirred model allows for prediction of MRT based on three readily available experimental parameters (fu_mic_, fu_p_, and CL_int_). While quantitative predictions of MRT based on *in vitro* data are still an active area of research, the categorical accuracy obtained using the approach presented in this work is a substantial improvement compared to the alternative method LipMetE.

The data shared regarding the use of mechanistic PK MPO scores in two small molecule drug discovery projects highlights the usefulness of these scoring functions in prioritizing compounds for *in vitro* and *in vivo* testing. In both Project 1 and 2, the mechanistic PK MPO approach recognizes the superior profile of the short-listed clinical candidates compared to the rest of the molecules synthesized during the project’s lifetime (AUCROC between 0.95 and 0.99); all molecules short-listed for clinical experiments were found in the top 10^th^ percentile. In addition, 5 out of 6 short-listed compounds were in the top 2^nd^ percentile. Similar to short-listed compounds, molecules tested in PK experiments are associated with significantly higher scores compared to molecules that did not progress towards *in vivo* testing (AUCROC = 0.86). Finally, in both Project 1 and 2, the mechanistic PK MPO scores increase chronologically, highlighting its value in tracking compound quality progression.

The combination of mechanistic PK MPO and IVIVc approaches presented in this work has the potential to significantly accelerate small molecule drug discovery campaigns and reduce *in vitro* and *in vivo* testing, as observed internally at during the Project 1 back-up effort.

## ASSOCIATED CONTENT

Supporting Information. The following files are available free of charge. Excel File with datasets used in this study.

## AUTHOR INFORMATION

Author Contributions:

FB had the idea.

FB, LJ, and JB developed method.

FB, DC, FL, VJ and LJ performed data analysis

FB, VJ, FB, and LJ wrote the manuscript

## ACKNOWLEDGEMENT

We thank Christoph Zapf, Natalie Hosea, Ivar Mcdonald, and Stephen Johnson for spending time reading manuscript and providing suggestions. Thanks to scientists at BMS who are involved in projects to generate data used for our analysis.

## ABBREVIATIONS

MPO: Multiparameter optimization
AUCROC: area under curve of receiver operating characteristic curve
AUC: area under the simulated PK curve
ML: machine learning
PK: pharmacokinetic
NME: new molecular entities
PBPK: physiologically based PK
ADME: absorption distribution metabolism elimination
DDI: drug-drug interaction
PD: pharmacodynamic
IVIVc: *in vitro* to *in vivo* correlation
MEC: minimum efficacious concentration
F: oral bioavailability
CL: clearance
fu_p_: fraction unbound in plasma
T_1/2_: half-life
VD_ss_: Volume of distribution
fu_mic_: fraction unbound in microsome
K_e_: rate of elimination
MRT: mean residence time
FN: fraction neutral
F_h_: fraction surviving first pass elimination in the liver
F_g_: fraction surviving first pass elimination in the gut following oral absorption
F_a_: fraction absorbed in the intestine
D_abs_: absorbable dose
P_eff_: effective jejunum permeability
P_app_: apparent permeability
k_a_: rate constant for absorption
fu_media_: binding estimates in potency media
TI: therapeutic index
ROC: receiver operating characteristic
fu_t_: fraction unbound in tissues
LipMetE: lipophilic metabolic efficiency

